# Simultaneous Deletion of Transient Receptor Potential Vanilloid 3 and Cacna1h Undermines CA^2+^ Homeostasis in Oocytes and Fertility in Mice

**DOI:** 10.1101/443622

**Authors:** Aujan Mehregan, Goli Ardestani, Ingrid Carvacho, Rafael Fissore

## Abstract

In mammals, calcium (Ca^2+^) influx fills the endoplasmic reticulum, from where Ca^2+^ is released following fertilization to induce egg activation. However, an incomplete index of the plasma membrane channels and their specific contributions that underlie this influx in oocytes and eggs led us to simultaneously knock out the transient receptor potential vanilloid, member 3 (TRPV3) channel and the T-type channel, Ca_V_3.2. Double knockout (dKO) females displayed subfertility and their oocytes and eggs showed significantly diminished Ca^2+^ store content and oscillations after fertilization compared to controls. We also found that the cell cycle stage during maturation determines the functional expression of channels whereby they show a distinct permeability to certain ions. In total, we demonstrate that TRPV3 and Ca_V_3.2 are required for initiating physiological oscillations and that Ca^2+^ influx dictates the periodicity of oscillations during fertilization. dKO gametes will be indispensable to identify the complete native channel currents present in mammalian eggs.

## INTRODUCTION

Mammalian egg activation is a widely researched field, as it is the first stage of embryo development. During this event, the egg is induced to undergo changes such as resuming and completing meiosis, remodeling its outer cortex to block polyspermy, reorganizing the cytoskeleton and meiotic spindle, undergoing pronuclear formation and DNA synthesis, and translating and changing maternal mRNA and protein levels to commence mitotic cycles (Horner and Wolfner, 2008; Florman and Fissore, 2015). These changes are commonly referred to as egg activation.

In this species, fertilization induces embryo development after the sperm fuses to a mature metaphase II (MII) oocyte (egg), and initiates a series of precise rises in the intracellular concentration of free calcium ([Ca^2+^]_i_), known as oscillations. The oscillations are ultimately responsible for triggering embryonic development via modification of proteins that regulate the resumption and completion of meiosis (Miyazaki and Igusa, 1981; Ducibella et al., 2002; Ozil et al., 2005). Ca^2+^ oscillations rely on Ca^2+^ influx from the extracellular media to replenish the stores (Igusa et al., 1983; Wakai and Fissore, 2013). Currently, the molecule(s)/channel(s) responsible for this influx has not yet been completely established.

During maturation – the process initiated following the surge of luteinizing hormone (LH) – and prior to ovulation and fertilization, the oocyte undergoes a plethora of changes including the increase of Ca^2+^ store content ([Ca^2+^]_ER_), which requires Ca^2+^ influx (reviewed in Wakai et al., 2011; Wakai et al., 2011; Whitaker, 2006). Previous studies have demonstrated that [Ca^2+^]_ER_ and Ca^2+^ influx are carefully regulated during maturation; while Ca^2+^ influx progressively decreases, [Ca^2+^]_ER_ content increases (reviewed in Wakai et al., 2011). This strict regulation is necessary because an excess of Ca^2+^ content and/or influx can predispose eggs or oocytes to parthenogenetic activation, fragmentation and/or apoptosis (Gordo et al., 2002; Ozil et al., 2005), whereas a deficit might impede cellular functions, including protein synthesis, completion of maturation, and initiation of embryonic development. Oocytes and eggs have several mechanisms to regulate changes in [Ca^2+^]_i,_ including pumps, channels, and exchangers; the PM Ca^2+^-ATPase (PMCA) and Na^+^/Ca^2+^ exchangers extrude excess Ca^2+^, while the sarco-endoplasmic reticulum Ca^2+^-ATPases reuptake Ca^2+^ into the ER thereby refilling its stores (reviewed in Berridge et al., 2000; Bootman et al., 2001; Wakai et al., 2011). This complement of molecules is known as the Ca^2+^ toolkit, one that every cell type possesses to regulate Ca^2+^ and trigger crucial processes such as muscle contraction, exocytosis, and metabolism, among others (Berridge et al., 2003).

The identification of the channels responsible for Ca^2+^ homeostasis in mammalian oocytes and eggs is largely incomplete. Among the plasma membrane (PM) channels, the mammalian transient receptor potential (TRP) family of channels include six subfamilies and nearly 30 human members that are expressed in multiple cell types and tissues (Wu et al., 2010). We have demonstrated the presence of two family members in oocytes and eggs including TRP Vanilloid, member 3 (TRPV3) (Carvacho et al., 2013). TRPV3 allows divalent cations such as strontium (Sr^2+^) and Ca^2+^ into cells and eggs, and importantly, it is essential for triggering parthenogenetic embryonic development using Sr^2+^ stimulation, though it is not required for normal fertility, as null females are fertile (Cheng et al., 2010; Carvacho et al., 2013). Another channel involved in Ca^2+^ homeostasis in oocytes and eggs is the T-type voltage-gated calcium channel, Ca_V_3.2 (Bernhardt et al., 2015), although *Cacna1h* null females are only mildly subfertile, which is consistent with the knowledge that changes in membrane potential during mouse fertilization are minor, and at the resting potential of oocytes and eggs a limited number of Ca_V_ channels are open (Igusa et al., 1983; Jaffe and Cross, 1984).

TRPV3 and Ca_V_3.2 channels are differentially expressed in oocytes and eggs. The functional expression of TRPV3 is nearly absent at the beginning of maturation at the germinal vesicle stage (GV), but rises steadily during maturation with its maximal expression being at the MII stage (Carvacho et al., 2013). On the other hand, the expression levels of Ca_V_ channels during oocyte maturation is unknown, although early electrophysiological recordings indicated that GV oocytes display greater current amplitude than ovulated eggs; however, the electrical properties of the channels expressed in GV oocytes were different from the protein expressed in eggs. Moreover, the total current was not corrected by cell area, thus it is not clear whether the increased expression observed of the channel at GV oocytes is accurate (Peres, 1986; Peres, 1987). Why these channels are differentially expressed and/or regulated during oocyte maturation requires further investigation.

Thus, despite identification of some channels in mammalian oocytes and eggs, the complete set of channels responsible for filling the internal Ca^2+^ stores and supporting oscillations has not been found. Furthermore, the ability to accurately probe the effects of channel inhibition on Ca^2+^ homeostasis in mouse eggs is hindered by the lack of specific and known pharmacological agents as well as by the lack of specific antibodies. Therefore, evaluation of Ca^2+^ store content, Ca^2+^ responses to agonists and fertilization in oocytes and eggs null for specific channel(s) is a necessary approach to identify the channel(s) that underlie Ca^2+^ homeostasis in these cells. In addition, these null oocytes and eggs will be an important platform to perform electrophysiological studies to identify the full complement of channel(s) as well as to assess the specificity of commonly used pharmacological inhibitors. To these ends, here we describe the generation of mice lacking both *Trpv3* and *Cacna1h,* and show that these females are subfertile compared to those lacking only one of the two channels. Most importantly, we found that oocytes and eggs of these dKO mice exhibit altered Ca^2+^ homeostasis and mount short-lived Ca^2+^ oscillations with reduced periodicity. Our findings therefore reveal insights into the Ca^2+^ channels required to initiate and maintain fertilization induced [Ca^2+^]_i_ oscillations in the mouse and possibly in other mammals.

## RESULTS

### Double null mice lacking Trpv3 and Cacna1h genes are subfertile

Our first goal was to generate a dKO mouse line lacking the *Trpv3* and *Cacna1h* genes. The rationale for this stemmed from the establishment of the single KO lines for these genes displaying little effect on Ca^2+^ homeostasis or influx in oocytes and eggs (Carvacho et al., 2013; Bernhardt et al., 2015). Besides examining how the absence of these channels would affect fertility, the simultaneous elimination of these channels would facilitate performing electrophysiological recordings to identify any remaining channel(s). Our ultimate goal is to pinpoint the channel(s) responsible for Ca^2+^ influx during oocyte maturation, fertilization and egg activation.

To obtain the dKO mouse line, single knockout mice were bred to generate the initial pool of double heterozygotes. Males and females of this generation were bred to generate the parent generation of dKO and wildtype (WT) mice that were used in the following studies. Germline deletion of the *Trpv3* and *Cacna1h* alleles was confirmed via PCR analysis using ear tissue DNA prepared from 21 day-old mice (Supplementary Fig. S1). We first investigated the possibility of obvious differences in ovarian size, ovulation rates, and on the rates of *in vitro* maturation. We found that there were no significant differences in ovarian weight and number of eggs ovulated post hormone stimulation between the groups (Supplementary Fig. S2A-B). Ovarian shape and size were also similar between the two groups (Supplementary Fig. S2C). It is worth noting that these evaluations were performed in young, 4-6-week-old, animals, which as shown in the subsequent figures, have fewer defects in fertility. There was also no delay in any stage of maturation in the dKO oocytes compared to rates observed in control oocytes when GV oocytes from WT and dKO mice were matured under *in vitro* conditions (Supplementary Fig. S2D). This data reinforces the notion that TRPV3 and Ca_V_3.2 channels are functionally present in mouse oocytes and eggs, but are not required for oocyte maturation or egg activation, at least in young female mice.

We then sought to evaluate the fertility of the females lacking the *Trpv3* and *Cacna1h* genes. The single knockout lines, *Trpv3^-/-^* and *Cacna1h^-/-^,* respectively, have previously been shown to be viable and fertile (Cheng et al., 2010; Chen et al., 2003; Carvacho et al., 2013; Bernhardt et al., 2015). WT mice were used as controls. Four females from each WT and dKO line were bred with five males of the same genotype for 36 weeks; *Trpv3*-knockout (V3KO) and *Cacna1h*-knockout (t-knockout; tKO) mating studies were performed with three pairs of mice.

Data from the first six litters was used for analysis. Our results show that the dKO line produced fewer numbers of pups, and the number of pups per litter decreased significantly by, or after, the third parturition (dKO: 5.58 ± 1.16 versus WT: 8.29 ± 0.62; p = 0.0115) (Fig. 1A). In contrast, V3KO females yielded a similar number of pups per litter compared to WT females; while tKO females yielded fewer pups per litter though non-significant compared to wildtype females (V3KO: 7.94 ± 1.7; tKO: 7.44 ± 0.98; p > 0.05 for both genotypes) (Fig. 1B). Similarly, the total number of pups yielded per female in each genotype after six parturitions was decreased by about 33% in the dKO line (dKO: 33.5 ± 10.7 versus WT: 49.8 ± 5.06, p = 0.037) (Fig. 1C). Lastly, we examined if there was a difference in the interval between litters. While dKO mice displayed a delay in this parameter, it remained statistically insignificant. It is also worth noting that after the third parturition in dKO females, and with each successive parturition, the number of neonatal deaths became prominent, with about 40-80% of pups dying per litter (data not shown). The total number of pups born from all females in each group varied significantly with a total of 191 pups yielded from dKO females versus 284 pups yielded from the controls (Supplementary Table S1). These results demonstrate that these channels are not required for fertilization nor to support embryo development to term, although they seem necessary for full fertility.

**Figure 1.**
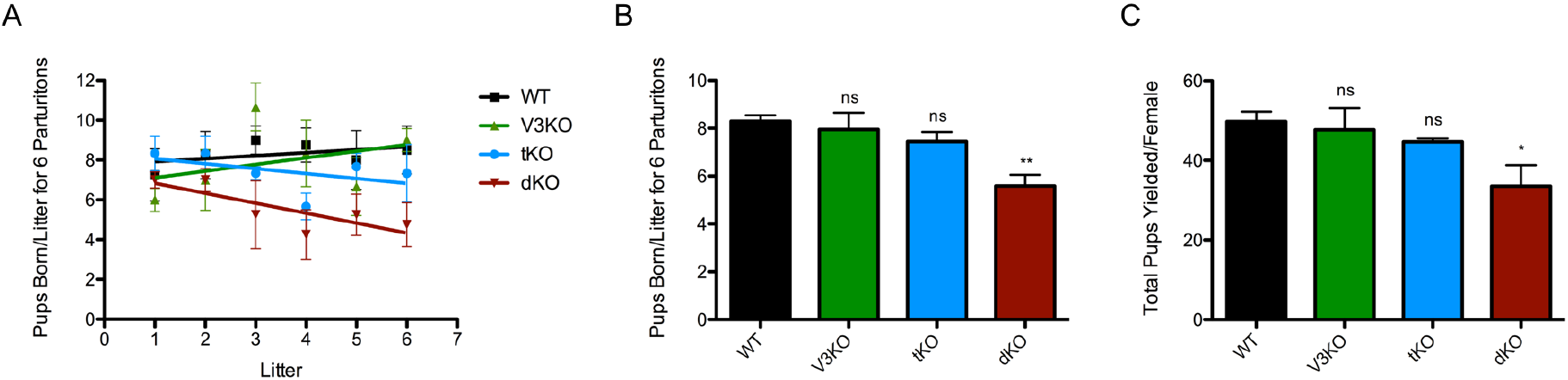
dKO females display subfertility. A-B: Number of pups born per litter for six parturitions. Linear regression in A was applied using data from each individual mating pair per genotype (WT n=4, V3KO n=3, tKO n=3, dKO n=4). Error bars represent standard error. B: Quantification of A, where mean ± S.E.M. for each genotype was as follows WT: 8.29 ± 0.62; V3KO: 7.94 ± 1.7; tKO: 7.44 ± 0.98; dKO: 5.58 ± 1.16. p (WT:dKO) = 0.0115 and p (WT:V3KO) or (WT:tKO) > 0.05. C: Total number of pups yielded per female. Mean ± S.E.M. for each genotype: WT: 49.8 ± 5.06; V3KO: 47.7 ± 9.45; tKO: 44.7 ± 1.53; dKO: 33.5 ± 10.7. p (WT:dKO) = 0.037 and p (WT:V3KO) or (WT:tKO) > 0.05. All means ± S.E.M. were calculated using column statistics in Prism GraphPad for each genotype.

### TRPV3 and Ca_V_3.2 currents are absent in dKO females

We used whole-cell patch clamp techniques to record and determine TRPV3 and T-type channel properties in MII eggs. Previous data established the expression of TRPV3 channels in mouse eggs and its potentiation by 2-Aminoethoxydiphenyl borate (2-APB) (Carvacho et al., 2013). 2-APB, besides being a nonspecific blocker of Ca^2+^ channels, was also identified as an inhibitor of IP_3_R1 (Maruyama et al., 1997). However, in eggs and at the concentration used here, 2-APB acts selectively on TRPV3 channels (Lee et al., 2016). In response to a voltage ramp (Fig. 2A), addition of 200μM 2-APB evoked an outwardly rectifying current with properties characteristic of TRPV3 (Hu, H. Z. et al., 2004), and congruent with Carvacho et al. (2013) (Fig. 2B). The current was present in WT eggs with a mean of 17.7 ± 4.2 pA/pF at +80 mV, but absent in dKO eggs (2.94 ± 1.5 pA/pF), which is comparable to basal currents. The inward current in both genotypes was statistically insignificant in the presence and absence of 2-APB. (Fig. 2B-C, respectively); thus, confirming the identity and absence of the channel. Next, we performed whole-cell patch clamp recordings to elicit Ca_V_3.2 currents. In response to a step protocol from -100 mV to +50 mV (Fig. 2A), we observed an I-V curve that agrees with T-type calcium channel activity. The peak of the currents in 20 mM extracellular Ca^2+^ was at -20 mV (Fig. 2D), which was consistent with previous reports (Bernhardt et al., 2015; Day et al., 1998; Peres, 1987). This current was absent in dKO eggs (WT: -2.81 ± 0.18 pA/pF versus dKO: 0.29 ± 0.01 pA/pF). To summarize, we show averaged current amplitudes at +80 mV and -80 mV (Fig. 2E), and at -20 mV (Fig. 2F). To confirm that dKO oocytes were truly null for both channels, we measured Ca_V_3.2 current, and subsequently, TRPV3 current in the same egg versus independent measurements in separate eggs, and observed absence of these currents in the dKO eggs (Supplementary Fig. S3).

**Figure 2.**
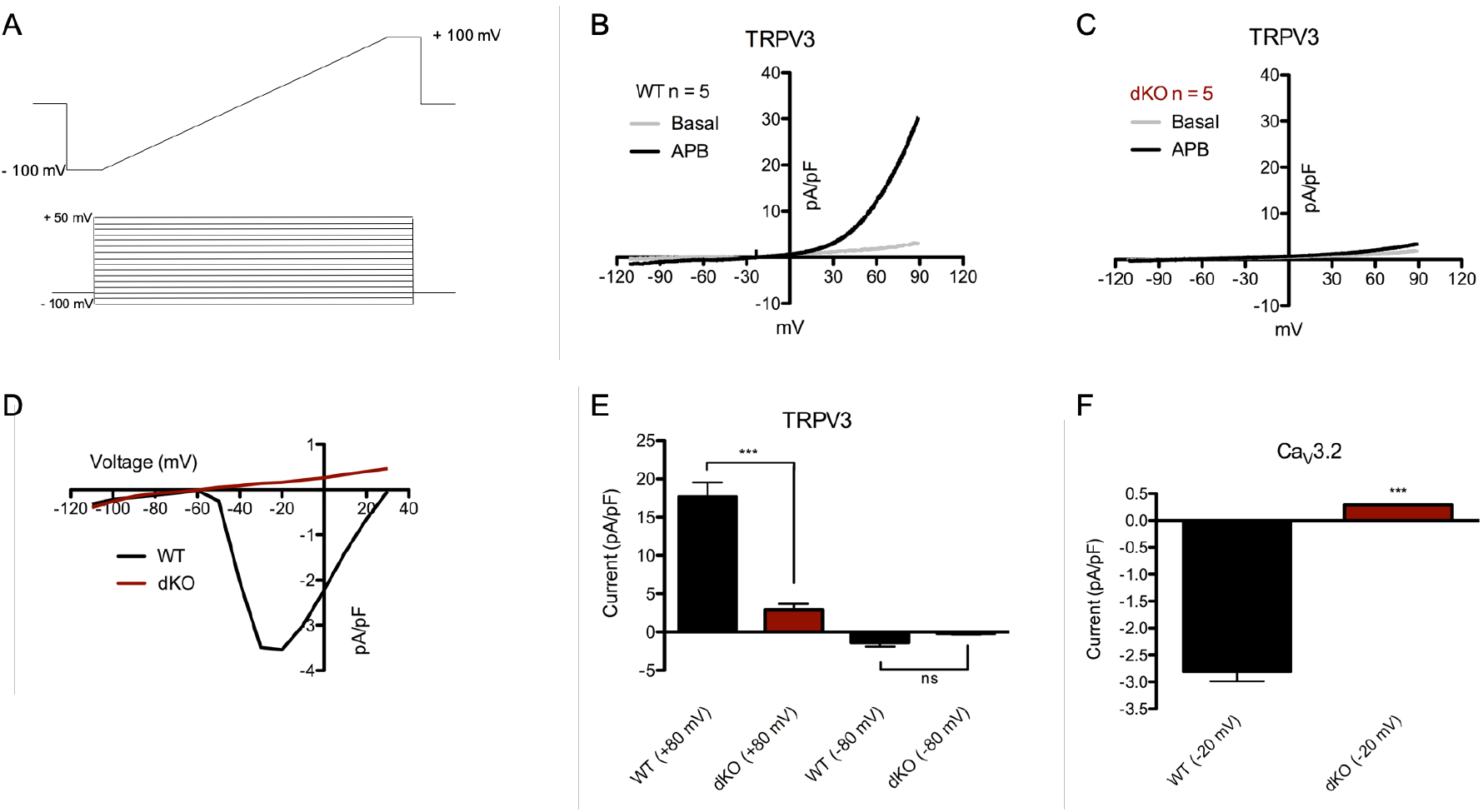
TRPV3 and Ca_V_3.2 currents are absent in dKO eggs. A, Top: ramp protocol from -100 mV to +100 mV to measure TRPV3 current (HP: 0 mV). Bottom: step protocol from -100 mV to +50 mV, every 10 mV to measure Ca_V_ channel activity (HP: -80 mV). B-C: Current-voltage (I-V) relationships in response to a ramp in the absence (grey trace) and presence (black trace) of 200 μM 2-APB. B: WT mean, n = 5 per trace. C: dKO mean, n = 5 per trace. D: I-V relationship in response to voltage step protocol. WT mean (black trace, n = 5) and dKO mean (red trace, n = 4). E: Averaged TRPV3 current responses in response to 200 μM 2-APB analyzed at +80 mV and -80 mV for WT (black bars; +80 mV: 17.7 ± 4.2 pA/pF; -80 mV: -1.38 ± 1 pA/pF) and dKO eggs (red bars; +80 mV: 2.94 ± 1.5 pA/pF (p < 0.0001); -80 mv: -0.198 ± 0.25 pA/pF (p = not significant)). F: Averaged Ca_V_3.2 current at -20 mV in WT eggs (−2.81 ± 0.18 pA/pF) versus dKO eggs (0.29 ± 0.01 pA/pF; p < 0.0001).

### Ca^2+^ stores are diminished in dKO females

The subfertility of the dKO mice suggested that the oocytes may have impaired Ca^2+^ homeostasis parameters. It has been previously documented that [Ca^2+^]_ER_ increases throughout maturation, which effectively plays a role in the preparation of the oocyte for fertilization (Jones et al., 1995; Wakai et al., 2011; Wakai and Fissore, 2013). Little is known about the mechanism by which oocytes accumulate Ca^2+^ in the stores during this process, although results from our laboratory suggest that the main source of increased [Ca^2+^]_ER_ is due to influx of external Ca^2+^ (Wakai et al., 2013). Ca_v_3.2 channels have also been shown to contribute to the increase in [Ca^2+^]_ER_ during oocyte maturation (Bernhardt et al., 2015), although the effects of TRPV3 channels were not examined. Nevertheless, given that TRPV3 and Ca_V_3.2 are important Ca^2+^ influx channels in oocytes, we hypothesized that the [Ca^2+^]_ER_ would be greatly diminished in dKO eggs.

To test this hypothesis, we directly examined in *in vivo* matured eggs the [Ca^2+^]_ER_ using Thapsigargin (TG), a sarcoendoplasmic reticulum Ca^2+^ ATPase (SERCA) inhibitor. SERCA is the pump that fills the ER, the major Ca^2+^ reservoir in the cell (Fig. 3A-B) (Jones et al., 1995; Kline and Kline, 1992; reviewed in Berridge, 2002). We observed a significant decrease in [Ca^2+^]_ER_ levels between the dKO (mean area of 1.72 ± 0.16, n = 18) and WT oocytes (mean area of 3.69 ± 0.22, n = 18; p < 0.0001); as estimated by quantification of the area under the curve (AUC) (Fig. 3C; left axis) and relative maximum amplitude (Fig. 3C; right axis). In the case of single channel KOs, while both parameters were also reduced, the reduction was only significant for V3KO oocytes (Fig. 3D-F). In the absence of extracellular Ca^2+^, the addition of TG partially empties the ER, and this promotes Ca^2+^ influx to refill the stores when extracellular Ca^2+^ is added back to the media. We tested the eggs’ ability to influx Ca^2+^ after TG by adding 2 mM CaCl_2_. Notably, there was no significant difference in Ca^2+^ influx capability between the WT and dKO groups (data not shown).

**Figure 3.**
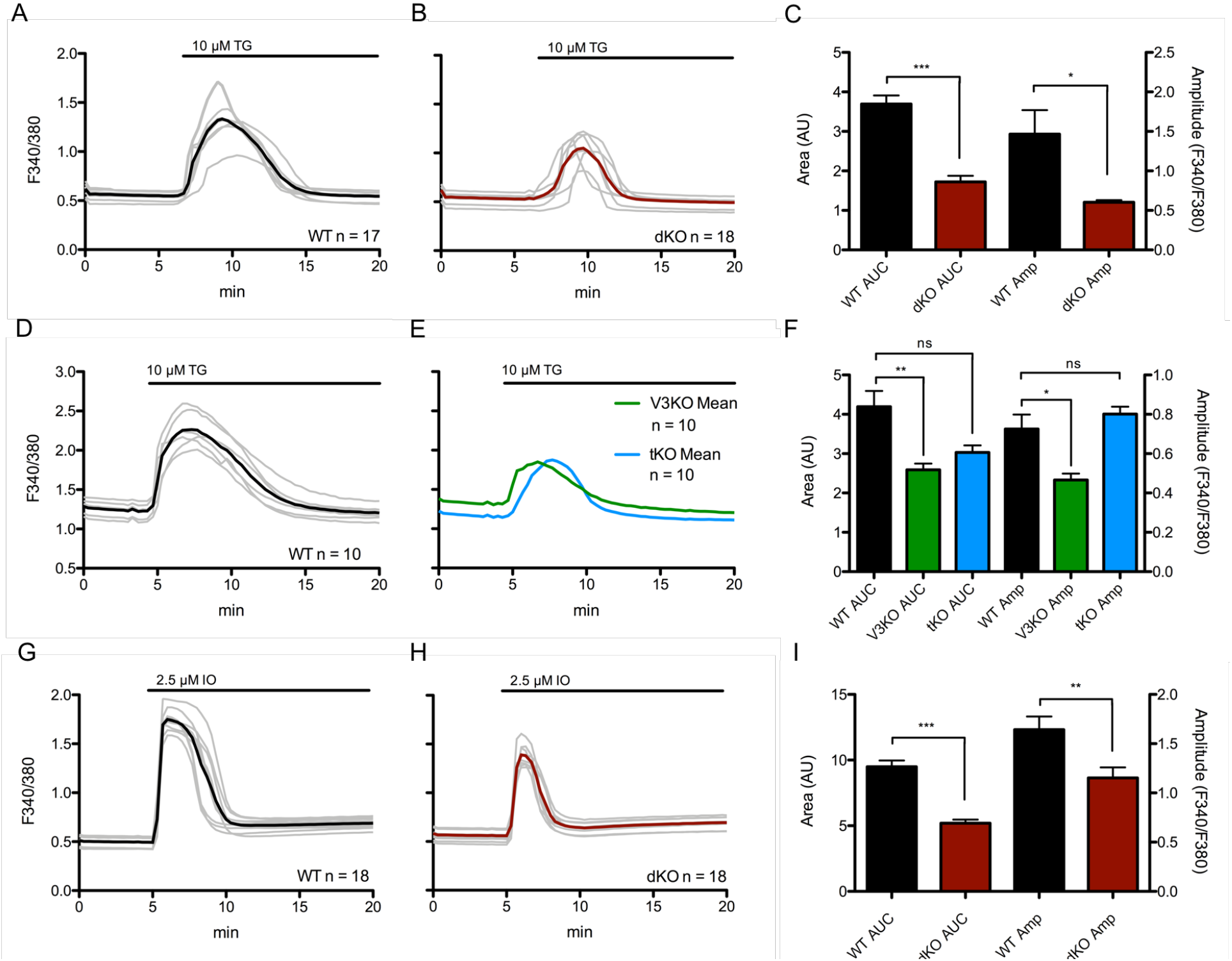
Ca^2+^ stores are reduced in dKO eggs. A-B: [Ca^2+^]_ER_ was measured by the addition of 10 μM thapsigargin (TG) under nominal Ca^2+^ conditions. TG added after 6 minutes. C: Summary of parameters measured. (Area under the curve (AUC) of WT: 3.69 ± 0.22, n = 17; dKO: 1.72 ± 0.16, n = 18; p < 0.0001; Amplitude (Amp) of WT: 1.47 ± 0.31; dKO: 0.60 ± 0.03; p = 0.014). D-E: [Ca^2+^]_ER_ measurements in WT, V3KO, and tKO MII eggs. TG added after 5 minutes. F: Summary of parameters measured. (AUC of WT: 4.2 ± 1.6, n = 15; V3KO: 2.6 ± 0.61, n = 15; tKO: 3.0 ± 0.62, n = 12; p(WT:V3KO) = 0.0014; p(WT:tKO) > 0.05. Amp of WT: 0.725 ± 0.29; V3KO: 0.466 ± 0.13; tKO: 0.801 ± 0.13; p(WT:V3KO) = 0.0002; p(WT:tKO) > 0.05. G-H: Total Ca^2+^ store content approximated by the addition of 2.5 μM ionomycin (IO) under nominal Ca^2+^ conditions. IO added after 7 minutes. I: Summary of parameters measured. (AUC of WT: 9.31 ± 0.54, n = 18; dKO: 5.29 ± 0.22, n = 18; p < 0.0001; Amp of WT: 1.16 ± 0.03, n = 18; dKO: 0.753 ± 0.06, n = 18; p < 0.0001). AUC and relative max amplitude were calculated using AUC analysis in Prism software after addition of TG or IO. Baseline was calculated from mean of y values from x = 0 to x = 5 min. Black trace represents mean values, grey traces represent individual responses.

Next, we used a Ca^2+^ ionophore, Ionomycin (IO), to empty all Ca^2+^ stores in the cell (Fig. 3D-E). By analyzing the same parameters as above, we observed that the total Ca^2+^ content, AUC, was decreased by almost half in dKO mice (5.29 ± 0.22, n = 18) compared to the control, WT mice (9.31 ± 0.54, n = 18; p < 0.0001) (Fig. 3F, left axis) as was the maximum amplitude (Fig. 3F, right axis). Collectively, these results suggest that oocytes null for two Ca^2+^ influx channels can still maintain, but to a lesser degree, [Ca^2+^]_ER_ levels during maturation and in MII eggs, and therefore that TRPV3 and Ca_V_3.2 channels are required to obtain a full amount of [Ca^2+^]_ER_.

### Sr^2+^ influx and 2-APB responses are abolished in dKO oocytes and eggs

Sr^2+^ is a useful method to induce artificial egg activation leading to parthenogenesis in rodent eggs. In MII eggs, Carvacho et al. demonstrated that TRPV3 channels mediate Sr^2+^ influx (2013). In a subsequent study, Carvacho et al. demonstrated that Sr^2+^ influx occurred through a different channel(s) at the GV stage, as oscillations persisted in V3KO GV oocytes (2016). We therefore tested if exposing WT and dKO GV oocytes and eggs to 10 mM SrCl_2_ induced oscillations. We found that Sr^2+^ failed to induce oscillations in dKO MII eggs, whereas the WT eggs showed robust responses (Fig. 4A-B). Further, when eggs were incubated in 10 mM SrCl_2_- containing media for two hours then washed into culture media and evaluated for egg activation, dKO eggs did not show any signs of activation such as extrusion of the 2^nd^ polar body, pronucleus formation or cleavage, whereas controls showed complete egg activation (data not shown). Another way to test for the absence of TRPV3 is by examining the response to 2-APB. Remarkably, 2-APB potentiates TRP Vanilloid channels, members 1-3, and is the most used activator of TRPV3 (Chung et al., 2004; Hu, H. Z. et al., 2004; Hu, H. et al., 2009). Here, we show that at 200 μM, 2-APB does not induce a Ca^2+^ rise in the dKO eggs, but it does in WT eggs (Fig. 4A-B). This data confirms and reinforces the finding that 2-APB induces a Ca^2+^ rise in eggs through the TRPV3 channel and that our dKO mice lack TRPV3.

**Figure 4.**
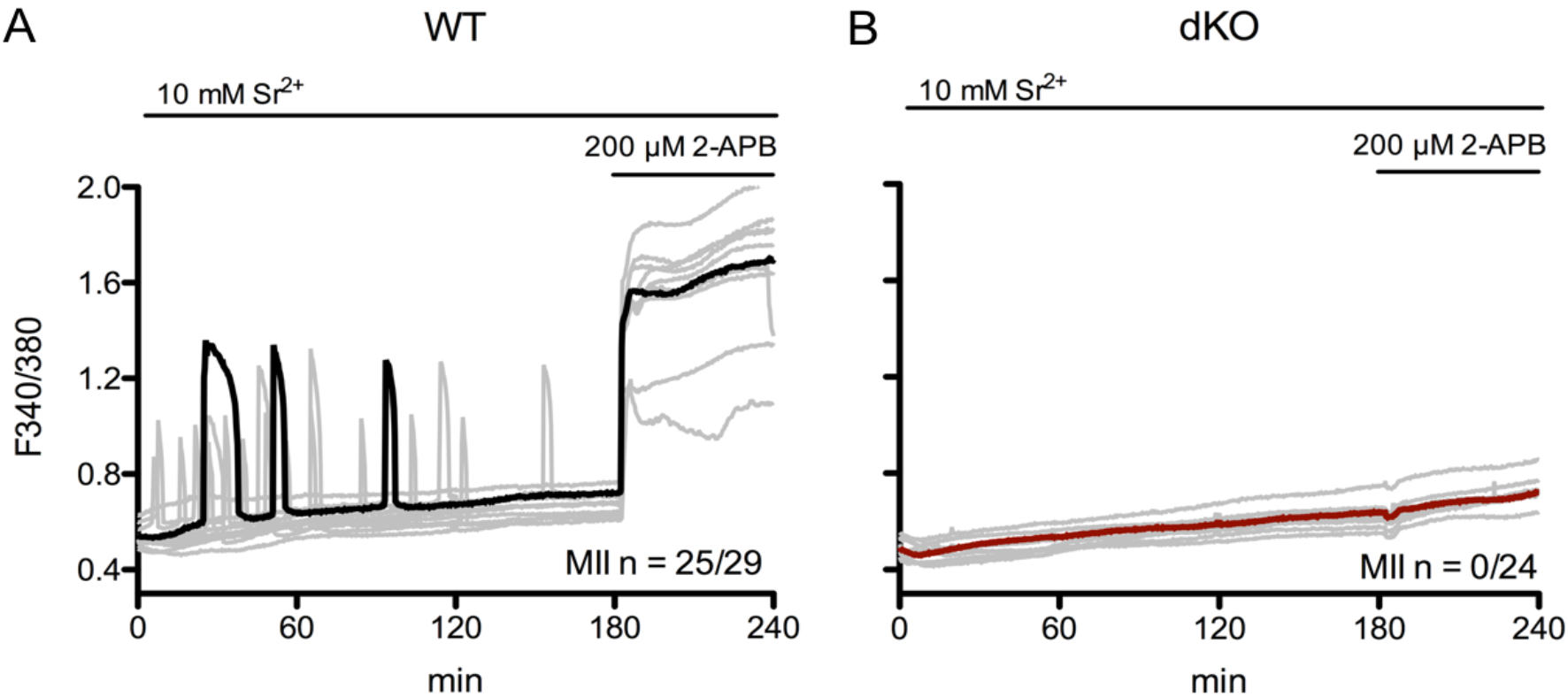
dKO eggs lacking TRPV3 and Ca_V_3.2 channels do not support Sr^2+^-induced oscillations. A-B: Oscillations induced in MII eggs by exposure to 10 mM SrCl_2_. A: Black trace shows representative WT egg displaying 3-4 oscillations in 60 minutes (n = 25/29) versus B: red trace, which shows no response (n = 0/24). 200 μM 2-APB was applied to media at the end of the experiment.

The absence of Ca_V_3.2 is harder to test without electrophysiology, as there are no specific agonists for these channels. Nevertheless, unpublished results from our laboratory suggested that Ca_V_3.2 may be an important mediator of Sr^2+^ influx at the GV stage. This is consistent with previously shown results where 10 mM SrCl_2_ exposure at the GV stage elicited spontaneous and irregular rises in WT and V3KO oocytes (Supplementary Fig. S4A; Carvacho et al., 2016). Importantly, we show here that in dKO oocytes, SrCl_2_ induced responses were largely absent (Supplementary Fig. S4B), demonstrating for the first time that Ca_V_3.2 channels underlie most of the Sr^2+^ influx in GV stage oocytes. Together, our results show that dKO mice lack functional expression of TRPV3 and Ca_V_3.2 channels and that their combined expression in oocytes ensures the influx of divalent cations throughout maturation.

### Ca^2+^ influx is diminished post-fertilization in dKO eggs

The introduction of PLCζ following sperm-egg fusion is thought to trigger the fertilization-associated [Ca^2+^]_i_ oscillations responsible for egg activation (Saunders et al., 2002). As a surrogate of fertilization, we tested the ability of the dKO eggs to mount Ca^2+^ oscillations following injection of PLCζ cRNA (Parrington et al., 1999; reviewed in Swann et al., 2006; Parrington et al., 2007). As shown, the time to initiation of oscillations was longer and the mean number of Ca^2+^ transients in the first 180 minutes was lower for dKO eggs (2.15 ± 0.18) versus control eggs (4.78 ± 0.17; p ≤ 0.0001) (Fig. 5A-C). tKO eggs displayed no significant difference in the frequency of oscillations (4.55 ± 0.55) compared to WT eggs (Fig. 5D). Furthermore, approximately only half of the dKO injected eggs mounted oscillations compared to 100% of the injected control eggs (Fig. 5A-C). Previous results with V3KO mice also showed responses comparable to controls (Carvacho et al., 2013). Collectively, we observed that the diminution of all parameters analyzed suggests that dKO eggs’ inability to influx the necessary amount of Ca^2+^ to support the filling and refilling of the ER undermines the ability to initiate and maintain timely oscillations (Fig. 5E-G).

**Figure 5.**
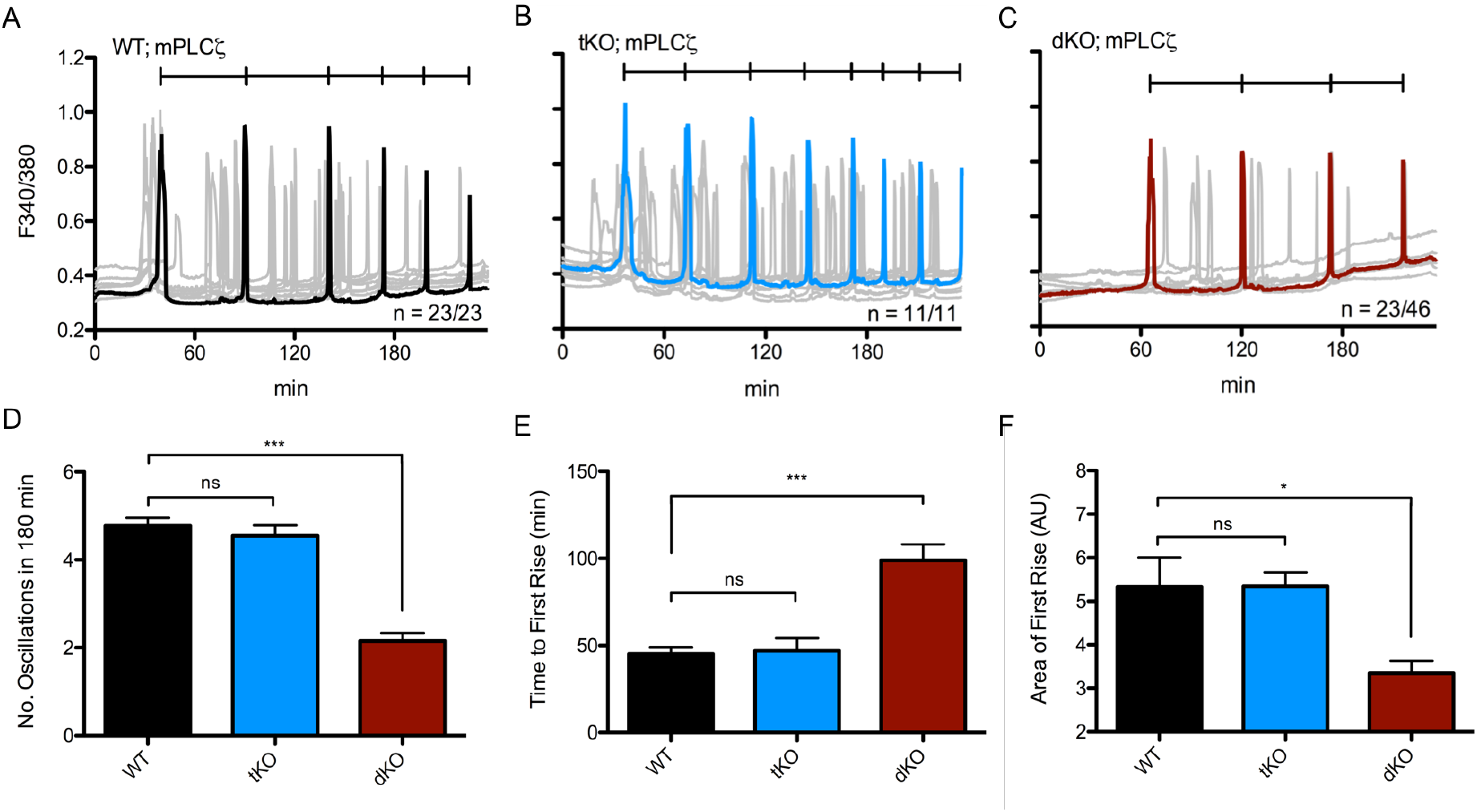
PLCζ cRNA-initiated oscillations are diminished post-PLCζ cRNA activation. Oscillations induced by microinjection of 0.01 μg/μL mouse PLCζ cRNA in WT eggs (A, n = 23) displaying 4.78 ± 0.17 oscillations in 180 minutes vs. tKO eggs (B, n = 11) displaying 4.55 ± 0.25 oscillations vs. dKO eggs (C, n = 46) displaying 2.15 ± 0.18 oscillations. Representative trace in black (A), blue (tKO), or red (dKO), individual traces in grey. D: Summary of parameters measured. p (WT:dKO) < 0.0001 and p (WT:tKO) was not significant. E: Time to reach first rise. x_0_ was start time of monitoring, x_f_ was 1^st^ point before inflection of first rise. p (WT:dKO) < 0.0001; p (WT:tKO) > 0.05. F: Area under the curve of the first rise. AUC was calculated via the integral of the first transient from x_1_ (first point of inflection) to x_2_ (last point before return to baseline). All measurements are represented as mean ± S.E.M. of every individual egg per genotype.

We next evaluated whether subsequent [Ca^2+^]_i_ rises in PLCζ cRNA injected eggs were comparable between dKO vs. WT eggs. We hypothesized that significant differences in certain parameters such as rise time and/or amplitude could suggest additional effects on the function of IP_3_R1s. To accomplish this, we examined the rise time of the third Ca^2+^ transient by measuring the slope between the first point of persistent increase in baseline Ca^2+^ until the value before reaching maximum amplitude (Deguchi et al., 2000). This parameter was not significant between groups (Supplementary Fig. S5A). Further, to rule out that the longer intervals in dKO eggs were due to reduced Ca^2+^ influx and not to the timing of injection or inability of dKO eggs to translate the cRNA, we quantified in WT and dKO eggs the fluorescent signal induced by injection of a cRNA encoding for a fluorescently tagged calcium-calmodulin kinase (CaMKII); this cRNA is expressed quickly and its accumulation does not cause cell cycle progression. We observed similar intensities at each time point in both groups (Supplementary Fig. S5B-C) indicating that dKO eggs are capable of efficiently translating injected cRNAs.

To extend the PLCζ cRNA results, we compared Ca^2+^ responses induced by fertilization using *in vitro* fertilization (IVF). As noted, TRPV3 channels appear unnecessary for the maintenance of fertilization-induced Ca^2+^ oscillations (Carvacho et al., 2013), and to a large extent a similar effect was observed for *Cacna1h^-/-^* mice (Bernhardt et al., 2015 and data in this manuscript). Nevertheless, we observed that WT eggs showed Ca^2+^ oscillations with normal frequency (Fig. 6A), whereas the frequency of oscillations in dKO eggs was substantially lower (Fig. 6B) with mean frequencies of 2.43 ± 0.14 oscillations per hour versus 0.667 ± 0.17 oscillations per hour, respectively (p ≤ 0.0001). We also quantified the stark difference in the area under the first transient, and observed that this parameter is significantly reduced in dKO eggs (7.92 ± 0.33 vs. 4.6 ± 0.68 AU) (Fig. 6C). The total number of Ca^2+^ transients was also decreased in the dKO eggs compared to the WT eggs (data not shown). Additionally, preliminary results following pre-implantation embryo development did not show any significant differences between the two groups (data not shown).

**Figure 6.**
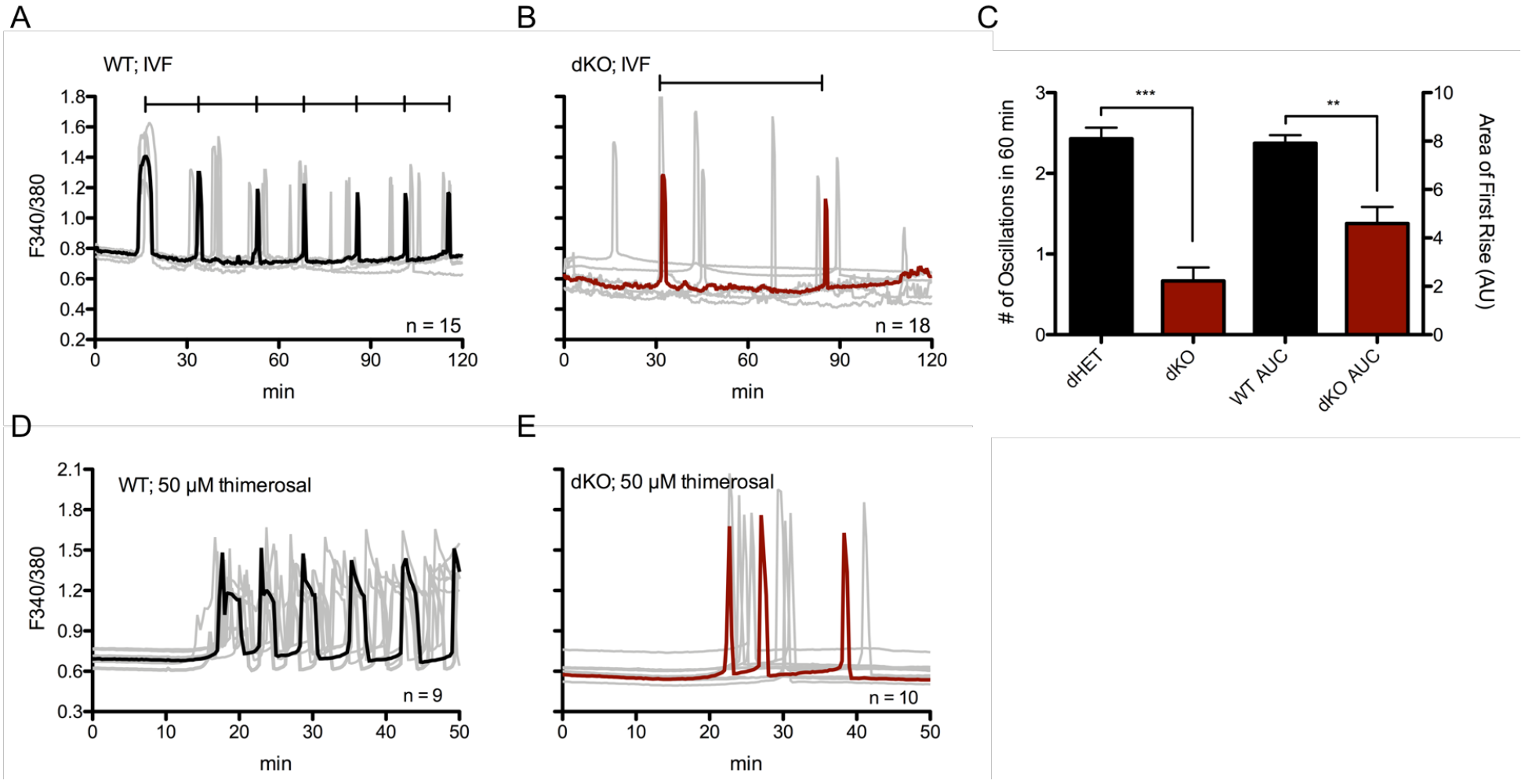
The absence of TRPV3 and Ca_V_3.2 channels significantly affect the pattern of Ca^2+^ oscillations post-fertilization or following addition of thimerosal. A: Oscillations induced by IVF in WT eggs (representative black trace with individual responses in grey traces; n = 15) who display 2.43 ± 0.14 oscillations in 60 minutes versus dKO eggs (B; n = 18) who display 0.667 ± 0.17 oscillations. C: Summary of parameters measured. Left y-axis: number of oscillations in 60 min, p < 0.0001. Right y-axis: area under the curve of first rise, p = 0.002. Bars represent mean ± S.E.M. of all traces per genotype. Statistical significance was calculated using two-tail t-test. D-E: 50 μM thimerosal induced fewer Ca^2+^ responses in dKO eggs (E, representative red trace, n = 10) vs. WT eggs (D, representative black trace, n = 9).

Lastly, we tested the relative sensitivity of dKO eggs to oscillate via chemical stimulation by monitoring WT and dKO eggs in media supplemented with 50 μM or 100μM thimerosal. Thimerosal is an oxidizing agent that is thought to enhance the sensitivity of calcium induced calcium release (CICR) and IP_3_Rs (Cheek et al., 1993; Swann, 1991), although its impact on PM Ca^2+^ channels was never tested. We show here that in response to thimerosal, dKO eggs mounted oscillations with irregular patterns and with lower frequency than controls, 3.2 ± 0.43 vs. 6.6 ± 0.45 rises in 60 min, respectively (Fig. 6D-E; p < 0.05). These data suggest that TRPV3 and Ca_V_3.2 channels are not required for the initiation of fertilization or chemically induced oscillations, but remarkably affect the periodicity of such oscillations, and are therefore physiological contributors to the [Ca^2+^]_i_ responses during mouse fertilization.

### A TRPM7-like channel is functional in eggs of dKO mice

The fact that the deletions of *Trpv3* and *Cacna1h* did not fully prevent the filling of [Ca^2+^]_ER_ and that fertilization induced oscillations were slowed but not prevented suggests the presence of Ca^2+^ influx by another channel(s). TRP Melastatin 7 (TRPM7) presence has been identified using electrophysiology (Carvacho et al., 2016), and it is functionally expressed in oocytes. This unique chanzyme is modulated at different levels by the divalent cation magnesium (Mg^2+^) (Bates-Withers et. al., 2011). For example, free intracellular Mg^2+^ affects the PM-associated domain, whereas Mg^2+^-ATP regulates the kinase domain, and high extracellular concentrations of Mg^2+^ affect channel permeability, effectively blocking the channel (Bates-Withers et al., 2011). Remarkably, the concentrations of Mg^2+^ in commonly used culture media, like HEPES-buffered Tyrode’s lactate solution (TL-HEPES), may be high enough to partially obstruct channels such as the TRPM7 channel (Ozil et al., 2017).

In support of a possible role of this channel, it has recently been shown that fertilization-induced embryo development in several species is increased in media with lower concentrations of Mg^2+^ (Herrick et al., 2015), and that sperm-initiated oscillations were increased when measurements were performed in the presence of low levels of extracellular Mg^2+^ (Ozil et al., 2017). To determine if indeed extracellular Mg^2+^ ([Mg^2+^]_o_) was affecting Ca^2+^ oscillations, we monitored oscillations after PLCζ cRNA injection in Mg^2+^-containing and Mg^2+^- free environments. It is worth noting, that changes in [Mg^2+^]_o_ concentrations in unfertilized WT and dKO eggs had no effect on the eggs’ basal Ca^2+^ levels (data not shown). Nevertheless, we observed that the absence of Mg^2+^ greatly increased the frequency of oscillations, and bringing Mg^2+^ to concentrations found in most commercial media slowed the oscillations, which in some cases ceased to continue (Fig. 7A-B). We observed the same effects in dKO GV oocytes after inducing spontaneous oscillations with SrCl_2_ (Supplementary Fig. S6A-B), which suggests that TRPM7-like channels are expressed in oocytes and eggs of dKO mice.

**Figure 7.**
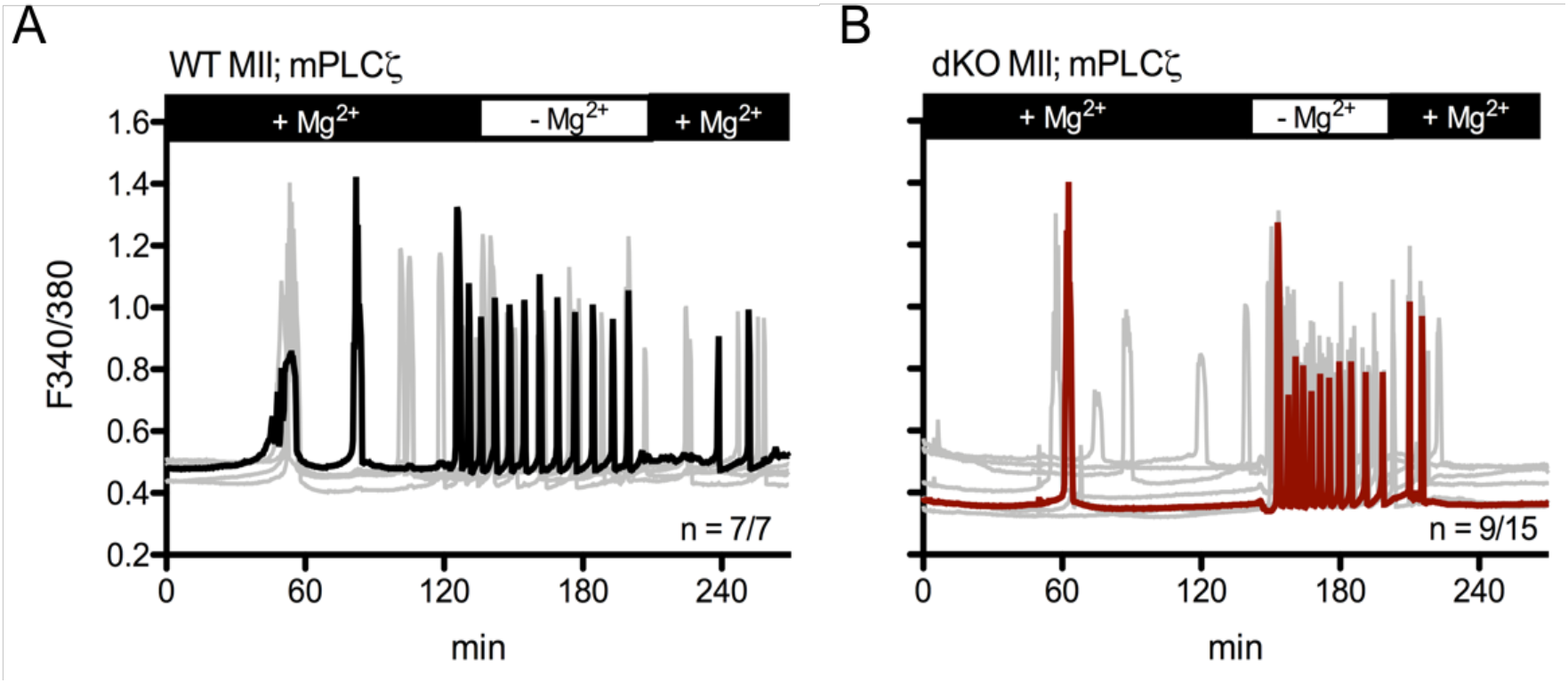
External Mg^2+^ modifies PLCζ induced Ca^2+^ oscillations in eggs. A-B: Oscillations induced after PLCζ cRNA microinjection in MII eggs. A: WT mean (black trace), n = 7/7. D: dKO mean (red trace), n = 9/15. Individual responses are shown as grey traces. Monitoring was continuous throughout changes in Mg^2+^ concentrations.

## DISCUSSION

Ca_V_3.2 channels were one of the first channels identified via molecular biology and electrophysiology in mouse eggs. Later, TRPV3 channels were identified using electrophysiology and KO animals (Peres, 1986; Peres, 1987; Kang et al., 2007; Bernhardt et al., 2015; Carvacho et al., 2013). Both channels are expressed in maturing oocytes, although definitive characterization of their expression and function during this process requires further investigation. Here, we studied the extent to which these channels are responsible for maintaining and increasing [Ca^2+^]_ER_ during maturation as well as their role in fertilization. Previously, it has been shown that mouse oocytes null for only TRPV3 (Carvacho et al., 2013) or only Ca_V_3.2 (Bernhardt et al., 2015) do not display major defects on fertilization or embryo developmental competency and are neither necessary nor sufficient for [Ca^2+^]_i_ oscillations. Nevertheless, given their prominent expression in oocytes and distinct expression patterns, we speculated their simultaneous elimination might have consequences in Ca^2+^ homeostasis and/or fertility. Our results show that their simultaneous absence greatly impacts Ca^2+^ homeostasis in oocytes and eggs and compromises the ability to initiate regularly spaced, frequent Ca^2+^ transients after fertilization. Nevertheless, while diminished, [Ca^2+^]_i_ oscillations are sustained, rendering dKO eggs a perfect platform to: 1) gain insights into the regulation of Ca^2+^ homeostasis during maturation and fertilization, and 2) assess the presence of other fundamental channels responsible for the totality of Ca^2+^ in oocytes and eggs.

Mouse oocytes and eggs contain a host of other potential sources of Ca^2+^ influx. Notably, another TRP family member, TRPM7, has been reported to be imperative for embryonic development (Jin et al., 2008). We recently demonstrated expression of TRPM7 in GV oocytes and MII eggs (Carvacho et al., 2016), though further experiments are required to clarify its function in oocytes and during pre-implantation development. It is nevertheless well known that TRPM7 is highly permeable to other divalent cations such as Zn^2+^ and Mg^2+^, and in fact Mg^2+^ homeostasis in the cell is largely mediated by TRPM7 (Bates-Withers et al., 2011). Thus, in addition to these ions, Ca^2+^ might also be permeating through this channel during development. Remarkably, [Mg^2+^]_o_ also acts as an antagonist of TRPV3 (Luo et al., 2012). Therefore, identification of the role of each channel would require studying Ca^2+^ responses in individual KO models as well as in models where several channels are eliminated.

### dKO Fertility

Ca_V_3.2 and TRPV3 channel function, singly or in combination, do not appear to be necessary for oocyte maturation, as a normal number of oocytes complete maturation and reach the MII stage in single KO females (Bernhardt et al., 2015; Carvacho et al., 2013), as well as our own results with dKO females in this study. Remarkably, we found a substantial decline in the fertility of dKO females, especially after the third litter, which also coincides with parturitions occurring at greater, though inconsistent, intervals. It is presently unclear what the underlying cellular or molecular reasons that progressively compromise fertility could be, as these defects are not observed in single KO lines. Our results show that the single KO is not enough to disrupt Ca^2+^ homeostasis, possibly because another channel(s) can effectively compensate. Simultaneous deletion, however, causes a significant effect, which might undermine embryo development. Future studies should examine histological sections of the ovaries at different ages, as well as collection of embryos following timed mating to elucidate the factor(s) compromising fertility in this model.

### Ca^2+^ Store Content in Eggs of dKO Mice

Using Ca^2+^-imaging measurements after addition of Ca^2+^ ionophore and/or TG, we found that dKO eggs showed vastly reduced [Ca^2+^]_ER_ store content over WT eggs. Moreover, oocytes and eggs from mice null for a single channel showed only minor effects on this parameter suggesting that these channel(s) could be compensating for each other’s absence. Importantly, the stores of dKO eggs were not empty, which suggests they are still capable of Ca^2+^ influx; as previously noted, TRPM7 is a candidate to mediate this influx. Further, the expression of TRPM7 might be augmented in dKO eggs, as suggested by the higher frequency of transients in the Mg^2+^-free experiments and the larger current in these eggs (data not shown).

### Sr^2+^ Responses in dKO Oocytes and Eggs

Sr^2+^ influx in MII eggs is mediated by TRPV3 (Carvacho et al., 2013), but not in GV oocytes, as Sr^2+^ oscillations are still observed in GV oocytes of V3KO mice (Carvacho et al., 2016). Research showed that Ca_V_ channels mediate divalent cation influx, including Sr^2+^, in cardiac Purkinje cells (Hirano et al., 1989a; Hirano et al., 1989b), and our results here with dKO mice confirm these results, as Sr^2+^ oscillations were greatly reduced in GV oocytes of dKO mice. We found that even in dKO GV oocytes, Sr^2+^ influx could be promoted if [Mg^2+^]_o_ was reduced or removed. Thus, TRPM7 might be the channel that mediates the residual influx of Sr^2+^ in dKO oocytes (Fig. 8). Additionally, our data suggests that the mechanism that favors Sr^2+^ influx changes during maturation, since dKO GV oocytes can still conduct some Sr^2+^, whereas MII dKO eggs cannot. Given that we show that Sr^2+^ mostly permeates through Ca_V_ channels in GV oocytes, and where this is not the case in MII eggs, our data suggests that during maturation Ca_V_ channels become progressively nonfunctional; how this is accomplished, though, may offer important insights into the regulation of Ca^2+^ homeostasis in oocytes.

**Figure 8.**
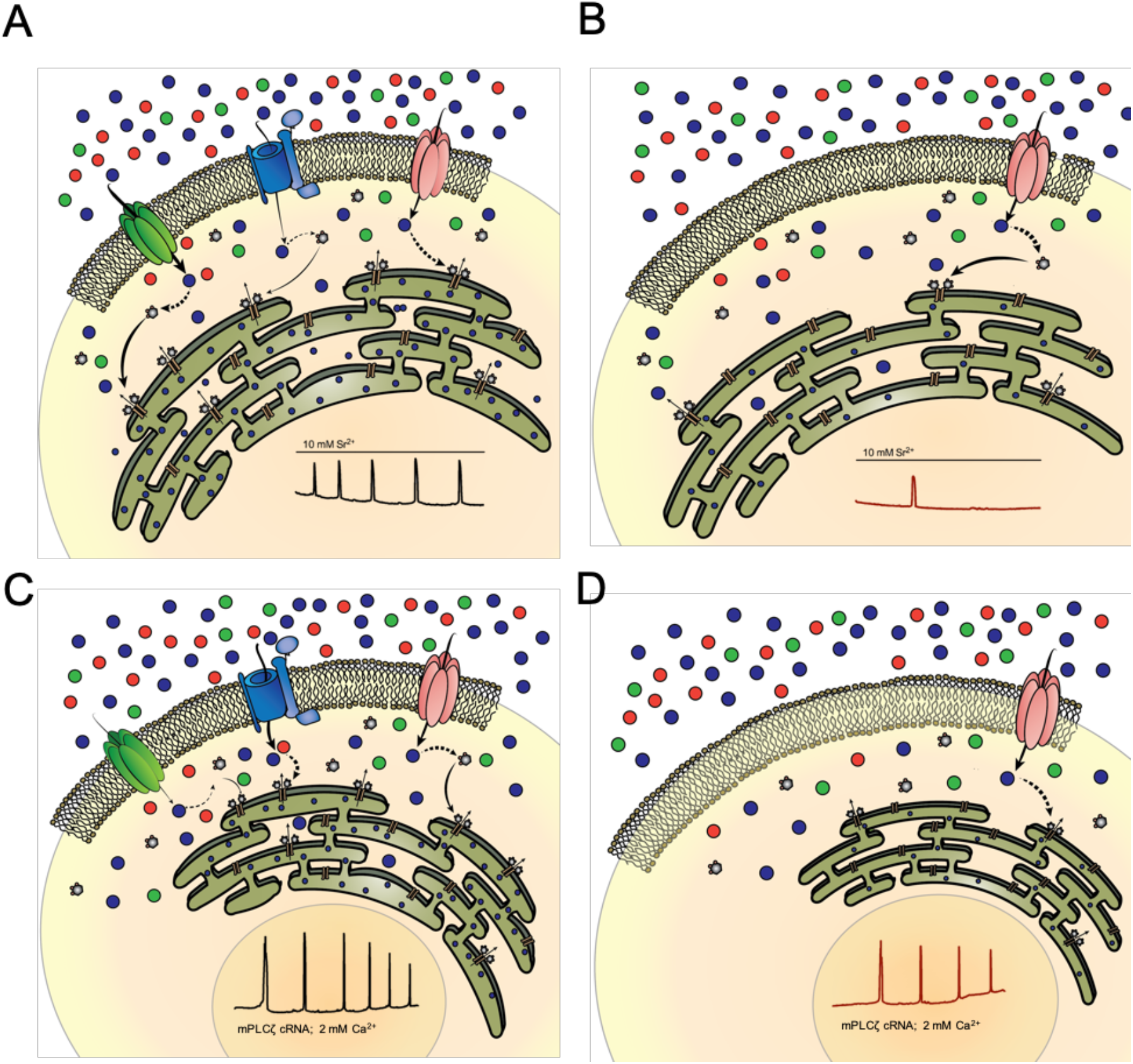
Divalent cation homeostasis is affected in dKO oocytes and eggs. A-B: Model GV oocyte expressing TRPV3 (green channel), Ca_V_3.2 (blue channel), and TRPM7 (red channel). A: WT oocyte shows normal divalent cation influx as indicated by weights of arrows, and normal pattern of SrCl_2_-induced oscillations. B: dKO GV oocyte exhibiting less Ca^2+^ influx, diminished pattern of SrCl_2_-induced oscillations, and thus decreased [Ca^2+^]_ER_. C-D: Model MII egg expressing TRPV3, Ca_V_3.2, and TRPM7. C: WT egg displays normal calcium homeostasis and mounts a regular pattern of fertilization-induced Ca^2+^ oscillations (black trace). D: dKO egg displays infrequent oscillations due to decreased Ca^2+^ influx (red trace). Dashed arrows represent degree of sensitization on a secondary messenger (IP_3_). Solid arrows represent direct translocation of secondary messenger. Ca^2+^ ions (blue), Sr^2+^ ions (orange) and Mg^2+^ ions (green).

### Ca^2+^ Oscillations Post-Activation and -Fertilization

Fewer dKO eggs showed Ca^2+^ oscillations in response to a variety of stimuli, and those that initiated oscillations showed a decreased frequency (Fig. 8). In all these cases, the persistence of the oscillations also seemed shortened. The mechanism whereby the absence of these channels undermines the mounting of robust [Ca^2+^]_i_ responses is unknown, but it might be that the reduced Ca^2+^ influx that causes slower refilling of the stores impairs the periodicity of the oscillations (Wakai et al., 2013). It has been proposed that intra-store Ca^2+^ levels sensitize the ER’s IP_3_R1s to the prevalent environmental [IP_3_], thus promoting Ca^2+^ release through this receptor (Taylor and Tovey, 2010). Therefore, the longer time needed to fill the stores to reach this threshold, the wider the intervals between rises.

Alternatively, the rate of refilling could influence the frequency of oscillations because the [Ca^2+^]_ER_level at which Ca^2+ “^leaks out” into the ooplasm, where it activates the catalytic activity of PLCζ (Sanders et al., 2018) leading to a quick increase in IP_3_ levels and [Ca^2+^]_i_ release, is slowed in the dKO eggs. This assertion seems to be supported by the finding that while the interval between Ca^2+^ rises is increased, the parameters of individual rises, for example the rate of rise of the third peak, does not appear different between WT and dKO eggs. These results suggest that the activation of PLCζ by the increasing concentrations of cytosolic Ca^2+^ is similar between WT and dKO eggs. Therefore, what we interpret to be different in dKO eggs is the slower influx of Ca^2+^ that delays the sensitization of IP_3_R1s and/or stimulation of PLCζ activity, in effect reducing the frequency of oscillations. Regardless of the specific mechanisms, our results are the first to show – without pharmacological manipulation, molecular overexpression, or abnormal concentrations of extracellular Ca^2+^ – that stimulated Ca^2+^ influx following sperm entry is a critical element in “setting the pace” of the Ca^2+^ oscillations during mammalian fertilization.

### Functional Role of TRPV3 and Ca_V_3.2 in Mouse Oocytes and Eggs

Given that oscillations persist in eggs of the dKO mice, a question that arises is why these channels are present in oocytes and eggs. In the case of Ca_V_3.2 channels, which are voltage-gated channels, the question is very relevant, as the mouse egg, a non-excitable cell, and in contrast to invertebrate species, experiences only a small change in membrane potential during fertilization (Jaffe and Cross, 1984; Igusa et al., 1983). Moreover, as mentioned, Ca_V_3.2-like currents have been measured in GV oocytes (Peres, 1986) and eggs (Peres, 1986; Day et al., 1998; Bernhardt et al., 2015); although at the reigning resting membrane potential in eggs of -30 to -40 mV (Peres, 1986), they are largely inactive. Nevertheless, a portion of the channels may display persistent inward currents at low voltages, referred to as “window currents” (Igusa et al., 1983). These currents have been detected in several cell types at or near to a membrane potential, comparable to the resting potential of unfertilized mouse oocytes and eggs (Bernhardt et al., 2015). Such a mechanism may be evident in the negative reversal potential observed in our dKO Ca_V_ recordings, which might reflect the lack of sufficient ions flowing across the membrane since the two main Ca^2+^ influx channels are missing, and the third purported channel, TRPM7, is blocked by high concentrations of Ca^2+^ (Li et al., 2006), in which our recordings are performed (see Materials and Methods).

The role of TRPV3 also needs some re-examination, since *Trpv3^-/-^* eggs do not show changes in oscillation frequency post-fertilization; elimination of both channels has devastating effects on the eggs’ Ca^2+^ store content and oscillations after fertilization. It is therefore plausible that eggs have redundant, compensatory channel(s) that sustain(s) normal oscillations in the event of the loss of a single channel. It is also possible that there are undetermined endogenous modulators of TRPV3 and Ca_V_3.2 that are present in oocytes and eggs, and how their activity is regulated will be the subject of future studies. Finally, although the exact function(s) of these channels remain(s) unknown, we provide evidence that they contribute to the maintenance of Ca^2+^ homeostasis pre- and post-fertilization. Gaining insight into the mechanism of Ca^2+^ influx during maturation and fertilization will aid in the generation of conditions that improve developmental competence especially of *in vitro* matured oocytes. Moreover, the identification of these channels as well as the development of specific channel blockers will contribute to the establishment of novel, non-hormonal methods of contraception to be used in humans, or to prevent the uncontrolled population growth of wild life species.

## MATERIALS & METHODS

### Animal Husbandry

WT and dKO mice were generated by breeding a female *Trpv3^-/-^* mouse (Cheng et al., 2010) (a generous gift from Dr. H. Xu, University of Michigan) with a mixed C57BL/6J and 129/SvEvTac background to a male *Cacna1h^-/-^* mouse (Jackson Laboratories, Bar Harbor, ME) with a B6;129-Cacna1h^tm1Kcam^/J background to generate F1 offspring heterozygous for *Trpv3 and Cacna1h* (dHET; +/-). Initial dKO and WT mice were obtained by intercross of dHETs and maintained on a mixed C57BL/6 and 129/SvEvTac background. Ear clips from offspring were collected prior to weaning, and confirmation of genotype was performed after most experiments.

### Oocyte Collection

Fully mature GV oocytes were collected from the ovaries of six-to-ten-week-old females that were superovulated by intraperitoneal (i.p.) injection of 5 IU pregnant mare serum gonadotropin (PMSG, Calbiochem, EMD Biosciences). GVs were collected and recovered into a HEPES-buffered Tyrode’s Lactate (TL-HEPES) solution supplemented with 5% heat-treated fetal calf serum (FCS, Gibco) and 100 μM IBMX to block spontaneous progression of meiosis. In-vivo matured metaphase-II (MII) eggs were collected by i.p. injection of 5 IU human chorionic gonadotropin (hCG, Calbiochem, EMD Biosciences) 46-48 hours post PMSG stimulation. Ovulated, MII-arrested eggs were obtained by rupturing the oviducts with fine forceps in TL-HEPES solution supplemented 5% FCS 12-14 hours post hCG stimulation. Cumulus cells were removed using 0.1% bovine testes hyaluronidase (Sigma, St. Louis, MO) and gentle aspiration through a pipette. All procedures were performed according to research animal protocols approved by the University of Massachusetts Institutional Animal Care and Use Committee.

### Genotyping/PCR Analysis

Mice were identified and genotyped using tissue from an ear clip, which was collected and lysed using tail lysis buffer (Tris pH 8.8 [50mM], EDTA pH 8 [1mM], Tween 20 [0.5%], proteinase K [0.3 mg/mL]). Genomic DNA was then stored at -20°C for later use in PCR analysis. Mouse genotyping was routinely performed using PCR analysis followed by fractionation on a 1.2% agarose gel. For *Trpv3*, F7622, 5’-GACATGCCATGCAAAAAACTACCA-3’ and R28432, 5’-GTCTGTTATATGTACAGGCATGG-3’ were used. The *Trpv3* WT and mutant alleles yielded products of 800 bp and 300 bp, respectively. For *Cacna1h*, 11395, 5’- ATTCAAGGGCTTCCACAGGGTA-3’, 11396, 5’-CATCTCAGGGCCTCTGGACCAC-3’, and oIMR2063, 5’-GCTAAAGCGCATGCTCCAGACTG-3’ were used. All primers were purchased from IDT Technologies (Coralville, IA). The *Cacna1h* WT and mutant alleles yielded products of 480 bp and 330 bp, respectively.

### Calcium [Ca^2+^]_i_ Imaging and Reagents

[Ca^2+^]_i_ monitoring was performed as previously reported by our laboratory (Kurokawa et al., 2007). Briefly, eggs were loaded with the Ca^2+^ sensitive dye Fura-2-acetoxymethyl ester (Fura-2AM, Molecular Probes; Invitrogen). Oocytes/eggs were loaded with 1.25μM Fura-2AM supplemented with 0.02% pluronic acid (Molecular Probes) for 20 min at room temperature. To estimate [Ca^2+^]_i_, oocytes/eggs were thoroughly washed and immobilized on a glass bottom monitoring dish (Mat-Tek Corp., Ashland, MA) submersed in FCS-free TL-HEPES under mineral oil. Oocytes/eggs were monitored under a Nikon Diaphot microscope outfitted for fluorescence measurements. The objective used was a 20X Nikon Fluor. The excitation lamp was a 75 W Xenon lamp, and emitted light >510 nm was collected by a cooled Photometrics SenSys CCD camera (Roper Scientific, Tucson, AZ) using NIS-Elements software (Nikon, Melville, NY). Oocytes/eggs were alternatively illuminated with 340 nm and 380 nm light by a MAC5000 filter wheel/shutter control box (Ludl Electronic Productions Ltd.), and fluorescence was captured every 20 s.

For experiments where the concentration of Mg^2+^ was changed, two blunt capillaries were secured to micromanipulators on either side of the glass bottom dish and inserted into the monitoring drop. One capillary was attached to a perfusion manifold via polyethylene tubing leading to open syringes filled with either normal, Mg^2+^ containing media, or Mg^2+^-free media. The other capillary was connected via polyethylene tubing to a closed syringe that acted as a manual vacuum when suction was applied with the plunger. Media was flowed in a slow, laminar fashion over the eggs during fluorescence intervals, while suction was simultaneously applied to change the concentration of Mg^2+^. Complete perfusion lasted about 2 min, and monitoring was continuous throughout the process.

For experiments using thimerosal, thimerosal (sodium ethylmercurithiosalicylate; Sigma) was prepared fresh daily by diluting it in TL-HEPES containing 2 mM CaCl_2_. Monitoring was performed in Ca^2+^-containing TL-HEPES without FCS and thimerosal was added after 5-7 min of baseline recording.

To examine the role of Ca^2+^ influx in refilling [Ca^2+^]_ER_, we monitored eggs in nominal Ca^2+^-free, FCS-free TL HEPES. After a 5-8-min baseline recording, [Ca^2+^]_ER_ levels were assessed by the addition of 10μM Thapsigargin (TG; Calbiochem, San Diego, CA), an inhibitor of the sarcoendoplasmic reticulum Ca^2+^-ATPase (SERCA) pump, which induced a Ca^2+^ leak via an unknown mechanism. TG-induced Ca^2+^ rises were regarded as [Ca^2+^]_ER_ content that could be estimated from the area under the curve of the [Ca^2+^]_i_ rise using Prism (GraphPad Software, La Jolla, CA). When [Ca^2+^]_i_ returned to near baseline values, ∼35 min after TG addition, 2-5 mM CaCl_2_ was added to the medium, and the amplitude of the [Ca^2+^]_i_ rise caused by the addition was used to estimate Ca^2+^ influx. In other experiments, the addition of 2.5μM Ionomycin (IO), a Ca^2+^ ionophore, was used to assess total store content of the egg. IO-induced Ca^2+^ rises were regarded as the total [Ca^2+^]_i_ that could be estimated from the area under the curve of the [Ca^2+^]_i_ rise using Prism.

### FRET and Calcium Imaging

To estimate the relative concentrations of Camui, the emissions of CFP, YFP and ratio imaging of the Camui (YFP/CFP) were monitored using a CFP excitation filter, dichroic beam splitter, CFP and YFP emission filters (Chroma technology, Rockingham, VT; ET436/20X, 89007bs, ET480/40m and ET535/30m). Eggs were then attached on glass-bottom dishes and placed on the stage of an inverted microscope. CFP and YFP intensities were collected every 20 second by a cooled Photometrics SenSys CCD camera and intensities compared between groups under examination. The rotation of excitation and emission filter wheels was controlled using the MAC5000 filter wheel/shutter control box (Ludl) and NIS-elements software. Imaging was performed on an inverted epifluorescence microscope using a 20x objective.

### Electrophysiology

Whole-cell currents were measured at 22-24°C using an Axopatch 200B amplifier digitized at 10 kHz (Digidata 1440A) and filtered at 5 kHz. Electrophysiology recordings were performed on the same day of egg isolation up to 8 h post-collection. Cumulus-free superovulated eggs were maintained in KSOM_AA_ at 37°C and 5% CO_2_. Shortly before measurement, eggs were aspirated briefly in acid Tyrode’s solution (pH 2.5) to remove the zona pellucidae. Data were analyzed using Clampfit (Molecular Devices) and Graphpad Prism. Pipettes of 1-3MΩ resistance were made from glass capillaries (593600, A-M Systems, CA), and typical seals of 1-4 MΩ were achieved before breaking into eggs. Series resistance was compensated by 40- 60%. The intracellular solution contained (in mM): 152 Cs-Methanesulfonate, 1 Cs-BAPTA, 10 HEPES, 2 MgATP, 0.3 NaGTP, 8 NaCl, pH: 7.3 adjusted with CsOH. The external solution contained (in mM): 125 NaCl, 6 KCl, 20 CaCl_2_, 1.2 MgCl_2_, 20 HEPES-NaOH, pH: 7.4. The response to 2-APB and mibefradil were measured in external solution containing (in mM): 140 NaCl, 10 HEPES, 10 glucose, 4 KCl, 1 MgCl_2_, 2 CaCl_2_. All voltages were corrected for calculated junction potentials present between the intracellular and external solutions before seal formation. TRPV3 currents were elicited by voltage ramps from -100 mV to 100 mV (600 ms, every 2 s), in the presence of 2-APB. The holding potential was 0 mV. Ca_V_3.2 currents were elicited by 50 ms duration depolarization steps from -100 mV to 50 mV in 10 mV increments. The holding potential was -80 mV. For experiments using inhibitors, seals were obtained in external solution containing 20 mM CaCl_2_ followed by equilibration in 2 mM CaCl_2_. Statistical analyses were performed using GraphPad Prism: t-test, paired, two-tailed p-value.

### Parthenogenetic Activation

For TRPV3-mediated egg activation, oocytes/eggs were collected as described above in TL-HEPES supplemented with 5% FCS (and 100 μM IBMX for GV experiments). For Ca^2+^ monitoring, oocytes/eggs were loaded with Fura-2AM, then immobilized to a glass-bottom monitoring dish (Mat-Tek Corp) under nominal Ca^2+^- and FCS-free TL-HEPES supplemented with 10mM SrCl_2_ (and 100 μM IBMX for GV experiments) submersed in mineral oil. For activation, eggs were incubated in 5% CO_2_ at 37° C for 2 h in Ca^2+^-free Chatot, Ziomek, or Bavister (CZB; Chatot et al., 1989) medium supplemented with either 3 mg/mL BSA or 0.01% polyvinyl alcohol (PVA), and 10mM SrCl_2_. Eggs were then washed and transferred to potassium-supplemented simplex optimized medium with amino acids (KSOM^AA^), and cultured to the 2-cell stage. Eggs were evaluated at 5-6 h and 22-24 h post treatment under phase contrast microscopy. Activated eggs were classified according to the following criteria: (1) PN group, consisted of zygotes forming a single PN with first and second polar bodies (5 h post-treatment); (2) cleaved group; eggs undergoing immediate cleavage after 24 h. Eggs without 2^nd^ polar bodies, PN formation, or those failing to cleave were considered as non-activated (MII egg). Fragmented eggs were excluded from analysis.

### Preparation of cRNAs and microinjections

The sequences encoding for Camui (generously gifted by Dr. Margaret Stratton, UMAss Amherst) and the full-length of mouse PLCζ cDNA, a kind gift from Dr. K. Fukami (Tokyo University of Pharmacy and Life Science, Japan) were subcloned into a pcDNA6 vector (pcDNA6/Myc-His B; Invitrogen, Carlsbad, CA). Plasmids were linearized with a restriction enzyme downstream of the insert and cDNAs were *in vitro* transcribed using the T7 or SP6 mMESSAGE mMACHINE Kit as previously described (Ambion, Austin, TX) according to the promoter present in the construct. A Poly (A)-tail was added to the mRNAs using a Tailing Kit (Ambion) and poly(A)-tailed RNAs were eluted with RNAase-free water and stored in aliquots at -80 °C. Microinjections were performed as described previously (Lee *et al.,* 2016). cRNAs were centrifuged, and the top 1–2 <u>μl</u> was used to prepare micro drops from which glass micropipettes were loaded by aspiration. cRNA were delivered into eggs by pneumatic pressure (PLI-100 picoinjector, Harvard Apparatus, Cambridge, MA). Each egg received 5–10 pl, which is approximately 1–3% of the total volume of the egg. Injected MII eggs were allowed for translation up to 4h in KSOM.

### Sperm Isolation

Spermatozoa for IVF procedures were obtained from 10-16-week-old male CD1 mice. The cauda epididymis of the sacrificed male was collected and sliced with scissors in 500μL of Toyoda, Yokoyama, Hosi (TYH) medium supplemented with 4 mg/mL bovine serum albumin (BSA; Sigma). The epididymis was incubated for 10-15 min at 37° C and 5% CO_2_ in TYH after which they were removed, whereas sperm were incubated for an additional 1 h under the same conditions.

### IVF

For standard IVF, expanded cumulus-oocyte-complexes were released from the oviduct and directly transferred to 90μL drops of TYH medium supplemented with 4 mg/mL BSA that was equilibrated overnight in 5% CO_2_ at 37° C, and 0.1-0.3 x 10^6^ sperm/mL were added. Complexes were incubated for 1 h, washed of excess sperm, and loaded with Fura-2AM for Ca^2+^ monitoring as described above. We also performed IVF in zona-free oocytes to detect [Ca^2+^]_i_ responses in WT and dKO mice. Procedures were performed as previously described (Bernhardt et al., 2015).

### Statistical Analysis

Values from three or more experiments performed on different batches of eggs were used for evaluation of statistical significance. Prism (GraphPad Software) was used to perform the Student’s *t*-test, one-way ANOVA, and graph productions. All data are presented as mean ± SEM. Differences were considered significant at *p* < 0.05 and denoted in bar graphs by the presence of asterisks.

### Chemical Reagents

Ionomycin, thapsigargin, PMSG, and hCG were purchased from Calbiochem (San Diego, CA). Fura-2AM and pluronic acid were purchased from Invitrogen (Carlsbad, CA). All other chemicals were from Sigma (St Louis, MO), unless otherwise specified.

## Acknowledgements

These studies were supported in part by NIH grants to R. A. F. (HD051872, HD092499). We would like to thank Ms. Changli He for technical support, Dr. James Chambers for electrophysiology and microscopy support, and Ms. Cristina Parrella for assistance with maintaining a breeding colony of mice and preliminary culture studies of pre-implantation embryos.

